# The *cis*-regulatory logic underlying abdominal Hox-mediated repression versus activation of regulatory elements in *Drosophila*

**DOI:** 10.1101/373308

**Authors:** Arya Zandvakili, Juli Uhl, Ian Campbell, Yuntao Charlie Song, Brian Gebelein

## Abstract

Hox genes encode a family of transcription factors that, despite having similar *in vitro* DNA binding preferences, regulate distinct genetic programs along the metazoan anterior-posterior axis. To better define mechanisms of Hox specificity, we compared and contrasted the ability of abdominal Hox factors to regulate two *cis*-regulatory elements within the *Drosophila* embryo. Both the Ultrabithorax (Ubx) and Abdominal-A (Abd-A) Hox factors form cooperative complexes with the Extradenticle (Exd) and Homothorax (Hth) transcription factors to repress the *distal-less* leg selector gene via the *DCRE*, whereas only Abd-A interacts with Exd and Hth on the *RhoA* element to activate a *rhomboid* serine protease gene that stimulates Epidermal Growth Factor secretion. By swapping binding sites between these elements, we found that the *RhoA* Exd/Hth/Hox site configuration that mediates Abd-A specific activation can also convey transcriptional repression by both Ubx and Abd-A when placed into the *DCRE,* but only in one orientation. We further show that the orientation and spacing of Hox sites relative to additional transcription factor binding sites within the *RhoA* and *DCRE* elements is critical to mediate appropriate cell- and segment-specific output. These results indicate that the interaction between Hox, Exd, and Hth neither determines activation vs repression specificity nor defines Ubx vs Abd-A specificity. Instead the precise integration of Hox sites with additional TF inputs is required for accurate transcriptional output. Taken together, these studies provide new insight into the mechanisms of Hox target and regulatory specificity as well as the constraints placed on regulatory elements to convey appropriate outputs.

**Author Summary:** The Hox genes encode a family of transcription factors that give cells within each region along the developing body plan a unique identity in animals from worms to mammals. Surprisingly, however, most of the Hox factors bind the same or highly similar DNA sequences. These findings raise a paradox: How can proteins that have highly similar DNA binding properties perform different functions in the animal by regulating different sets of target genes? In this study, we address this question by studying how two Hox factors regulate the expression of target genes that specify leg development and the making of liver-like cells in the developing fly. By comparing and contrasting how Hox target genes are activated and/or repressed, we found that the same Hox binding sites can mediate either activation or repression in a manner that depends upon context. In addition, we found that a Hox binding site that is normally regulated by only one Hox factor, can also be used by more than one Hox factor swapped into another target gene. These findings indicate that the specificity of a Hox factor to regulate target genes does not rely solely upon DNA binding specificity but also requires regulatory specificity.

## Introduction

Hox genes encode a family of highly conserved homeodomain transcription factors that specify distinct cell fates along the developing anterior-posterior (A-P) axis of the embryo [1]. Most animals contain at least five Hox genes that are often found clustered in the genome [2]. For example, *Drosophila melanogaster* encodes a single set of eight Hox genes that are split into two clusters (five in the *Antennapedia* complex and three in the *Bithorax* complex), whereas mammalian genomes have undergone Hox gene and cluster duplication resulting in four clusters that encode a total of 39 Hox genes [2]. While the number of Hox genes varies between animals, Hox genes share in common the property of instructing cells to adopt a "regional" (or "segment") identity within the organism by regulating the expression of downstream target genes [3]. Recent genomic studies have indicated Hox factors affect the expression of hundreds of downstream target genes [4-7]. Since each region or segment under the control of a specific Hox factor is composed of many cell- and tissue-types, these findings present two fundamental challenges in understanding how Hox genes sculpt the body plan: First, what makes one Hox factor different from another to specify distinct embryonic regions during development? Second, how can a regionally expressed Hox factor regulate downstream target genes in a cell- or tissue-specific manner?

Much of the focus on how Hox factors regulate distinct cell fates has been to define the mechanisms underlying target DNA binding specificity. Comparative studies between Hox factors revealed each binds highly similar AT-rich DNA sequences [8-10]. These findings raised a paradox: how can a family of transcription factors that bind nearly identical DNA sequences *in vitro* regulate distinct target genes and cell fates *in vivo*? A partial explanation for this phenomenon is that Hox factors form transcription factor complexes with additional proteins. The Extradenticle (Exd, *Drosophila*)/Pbx (vertebrate) and Homothorax (Hth, *Drosophila*)/Meis (vertebrate) families of transcription factors represent the best characterized Hox co-factor proteins [11-15]. Exd/Pbx and Hth/Meis proteins are widely expressed during development and bind DNA in a cooperative manner with Hox factors as well as with each other. As each protein in the Hox/Exd/Hth complex can bind DNA in a sequence-specific manner, Hox/Exd/Hth complexes enhance both target affinity and specificity [16-18]. Moreover, a comprehensive DNA selection assay revealed Hox factors gain discriminatory power when binding DNA with Exd, a concept called latent specificity [19]. Perhaps the best studied example of latent specificity is the regulation of the *Forkhead* (*Fkh*) gene by the Sex combs reduced (Scr) Hox factor during salivary gland development [20, 21]. A key feature of the Exd/Hox DNA binding site within the *Fkh* sequence is the presence of a narrow minor groove that is cooperatively bound by Scr only when in complex with Exd. Importantly, changing the *Fkh* DNA sequence to match a generic Exd/Hox consensus site (*Fkhcon*) resulted in a loss in Hox specificity as evidenced by ectopic reporter regulation via other Hox factors [22, 23]. Similarly, Crocker et al found that suboptimal Hox/Exd sites in the *shavenbaby* (*svb*) enhancer are regulated by two Hox factors (Ultrabithorax, Ubx and Abdominal-A, Abd-A), whereas "improving" these sites by more closely matching consensus binding sites resulted in a loss of specificity with many Hox factors activating target gene expression [24]. Thus, the formation of Hox-specific complexes on DNA with Exd/Pbx and/or Hth/Meis is a key mechanism that can yield Hox target specificity.

While the formation of Hox complexes with additional transcription factors can enhance DNA binding specificity, additional studies suggest Hox factors can differ in their regulatory potential once bound to DNA. For example, studies using the *Fkh* consensus element (*Fkh-con*) that provides generic Hox binding revealed that not all Hox factors that bound the element activated transcription as a subset instead repressed transcription [22]. Moreover, studies on additional *cis*-regulatory modules (CRMs) revealed that the Abd-A Hox factor can either activate or repress target gene expression when in complex with the Exd and Hth proteins. The *RhoBAD* regulatory element contains a highly conserved sequence (*RhoA*) encoding an adjacent set of Exd/Hth/Hox sites that recruits an Abd-A complex to mediate *rhomboid* (*rho*) activation in a subset of sensory organ precursor (SOP) cells [25-28]. The activation of *rho*, which encodes a serine protease that triggers the release of an EGF ligand, results in the induction of neighboring cells to form an essential set of hepatocyte-like cells known as oenocytes [29-32]. In contrast, the *Distal-less Conserved Regulatory Element* (*DCRE*) contains three Hox/co-factor binding sites that recruit Abd-A/Exd/Hth complexes to repress *Distal-less* (*Dll*) gene expression in the abdominal ectoderm [33-35]. *Dll*, which is an appendage selector gene that promotes leg formation in thoracic segments, is thereby restricted from the abdomen to block appendage formation in these segments [36]. Intriguingly, the *RhoA* and *DCRE* elements also differ in Hox specificity as the *DCRE* is regulated by both Abd-A and Ubx, whereas *RhoA* is regulated by only Abd-A. Thus, the studies of the *DCRE* and *RhoA* reveal that Hox factors can differ in both their target specificity and in their regulatory potential once bound to DNA.

What determines if an element is activated or repressed by a specific Hox factor? Current models suggest that the *RhoA* and *DCRE* CRMs integrate additional transcription factor inputs that dictate the sign of transcription. For example, *RhoA* requires a nearby Pax2 transcription factor binding site to mediate activation whereas the DCRE contains a nearby FoxG binding site (Sloppy-paired 1 (Slp1) and Slp2 are largely redundant *Drosophila* FoxG (FoxG) proteins) to mediate repression [26, 34]. How these factors are integrated with the specific Hox transcription factor complexes remain relatively unknown. In this study, we use a series of quantitative reporter assays to define the underlying *cis*-regulatory logic and mechanisms of Hox regulatory specificity by comparing and contrasting the ability of abdominal Hox factors to affect the *DCRE* and *RhoA* CRMs in conjunction with FoxG and Pax2.

## Results

### The *DCRE* mediates short-range transcriptional repression

The *DMX cis*-regulatory element activates *Dll* expression in the embryonic leg primordia [34, 36]. Prior studies revealed the *DMX* can be divided into two parts: the *DMEact* that conveys transcriptional activation in thoracic and abdominal segments and the *DCRE* that binds the Ultrabithorax (Ubx) and Abdominal-A (Abd-A) Hox factors to repress gene activation in abdominal segments [33, 37]. However, recent findings revealed abdominal Hox factors also repress the *DMEact* independent of the *DCRE* via unknown mechanisms [35]. To better define the *cis*-regulatory logic utilized by the *DCRE* to mediate abdominal repression, we used an assay that isolates the *DCRE* from the *DMEact* by placing it adjacent to three copies of the Grainyhead binding element (*3xGBE*). *3xGBE* sites are sufficient to activate gene expression throughout the ectoderm of the *Drosophila* embryo [35, 38]. Comparisons between *3xGBE-lacZ* (*G-lacZ*) and *3xGBE-DCRE-lacZ* (*GD-lacZ*) transgenes inserted into an identical chromosomal locus revealed the *DCRE* mediates robust repression in abdominal cells that co-express a *Drosophila* FoxG (Slp2) transcription factor (**Fig 1A-C**). To quantify abdominal repression, we measured β-gal intensity in Slp2+ cells and found that *G-lacZ* drives equivalent reporter levels in thoracic and abdominal segments whereas *GD-lacZ* embryos had ~70% less activity in abdominal segments relative to thoracic segments (compare **Fig 1B’’’** with **1C’’’**). Hence, the *GD-lacZ* assay provides a means to isolate and study *DCRE*-mediated repression independent from the more complex *DMX* element.

**Figure 1.**
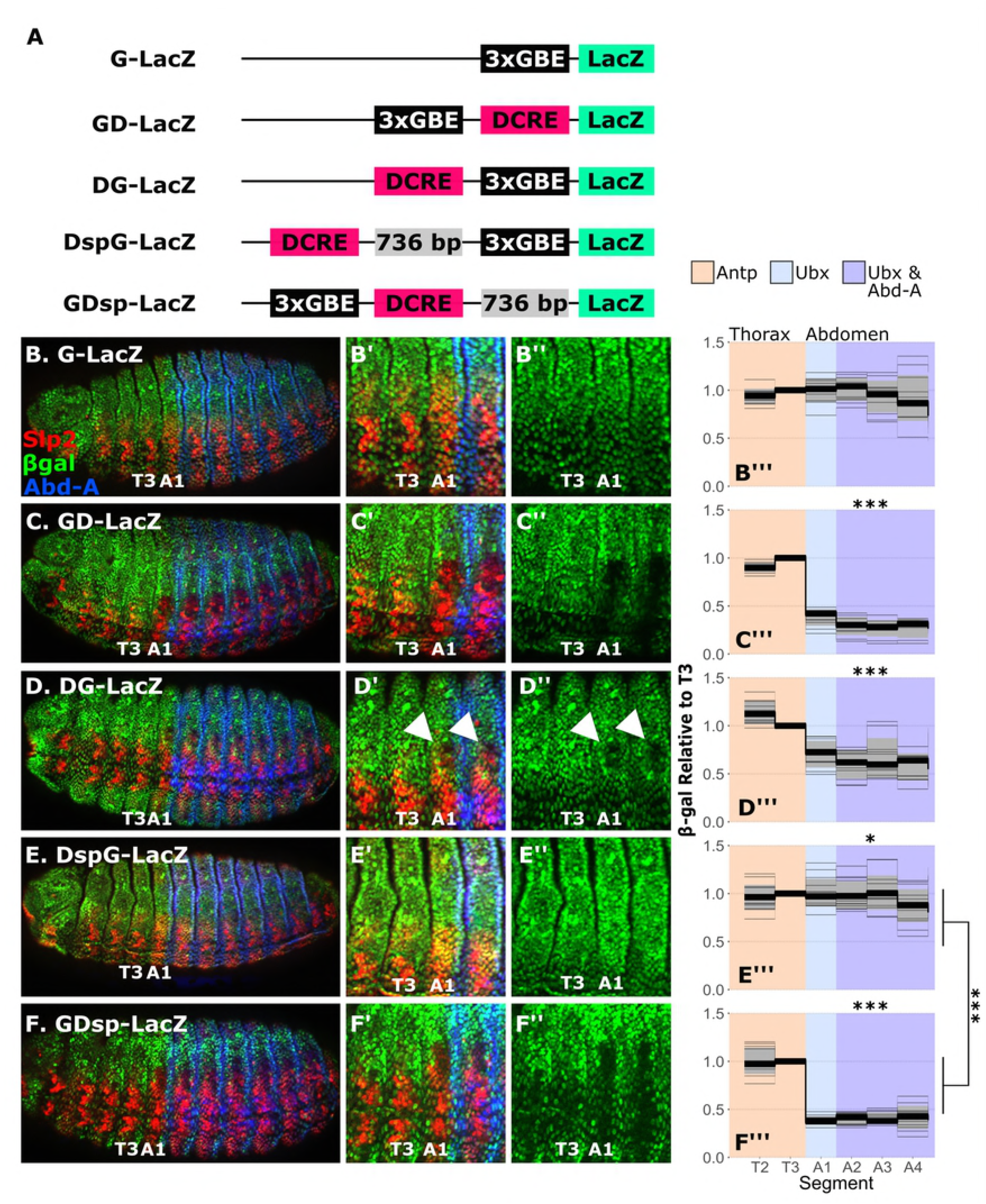
The *DCRE* is a short-range transcriptional repressor. **(A)** *LacZ* reporter constructs used to test *DCRE* activity. G = *3xGBE*; D = *DCRE*, sp = 736bp spacer from the *kanamycin* gene that is transcriptionally inert (Fig S1). **(B-F)** Lateral views of Stage 15 embryos expressing the indicated *LacZ* reporter constructs. Embryos were immunostained for β-gal (green), Slp2 (red), and Abd-A (blue). **(B’-F’)** High power view of the T2-A2 segments of the embryos in panels B-F. **(B’’-F’’)** Same as (B’-F’) but only showing β-gal stain**. (B’’’-F’’’)** Quantification of β-gal immunostain intensity among Slp2+ cells in segments T2-A4 relative to intensity in T3 segment. Background color indicates the Hox expression domain as shown by the key at top. Light gray lines are intensity values from individual embryos, while dark black line display median value between embryos. The grey ribbon displays mean intensity value +/- the standard deviation between embryos. Statistical comparison of mean abdominal β-gal intensity between reporter constructs was conducted using ANOVA with post-hoc Tukey’s test. Asterisks indicate significant difference from *G-LacZ* (“n.s.” = not significant; “*” p < 0.05; “**" p < 0.01; and “***" p < 0.001).

To define the range of repression activity of the *DCRE*, we engineered a series of constructs that alter the location of the *DCRE* relative to the *3xGBE* activation element (**Fig 1A**). First, we swapped the order of the *DCRE* and *3xGBE* (*DCRE-3xGBE-lacZ*, *DG-lacZ*) and found that in this configuration the *DCRE* mediates transcriptional repression, albeit weaker and predominantly in a subset of Slp2+ abdominal cells (**Fig 1D**). Next, we moved the *DCRE* further from the *3xGBE* (*DCRE-sp-3xGBE-lacZ, DspG-lacZ*) by inserting a 736 bp sequence from the *kanamycin* gene which was previously found to be transcriptionally inert [39]. Consistent with this spacer DNA not having significant transcriptional activity, we found that inserting it adjacent to the *3xGBE* did not significantly alter reporter expression (**Fig S1**). Importantly, the *DCRE* was unable to convey abdominal repression when it was separated from the *3xGBE* sites by the spacer DNA sequence (**Fig 1E**). However, moving the *3xGBE* adjacent to the distant *DCRE* (*3xGBE-DCRE-sp-LacZ, DGsp-lacZ*) rescued repression (**Fig 1F**). Thus, the *DCRE* functions as a relatively short-range element that represses transcription when placed adjacent to activation elements.

### Hox specificity and the *cis*-regulatory logic of Abd-A mediated *RhoA* activation versus *DCRE* repression

The *Ubx* and *abd-A* Hox genes encode nearly identical homeodomains (55 of 60 amino acids), bind highly similar DNA sequences as monomers, and form similar transcription factor complexes with the Exd and Hth Hox co-factor proteins on DNA *in vitro* [8, 19]. Consistent with these findings, prior studies demonstrated that Ubx and Abd-A repress *Dll* via the *DCRE* [33, 34]. To determine if both Ubx and Abd-A repress the *DCRE* in the *GD-lacZ* assay, we first analyzed *GD-lacZ* embryos and found that only Ubx is expressed in Slp2+ cells of the first abdominal segment (A1) [40], whereas both Ubx and Abd-A are detected in Slp2+ cells in subsequent abdominal segments (**Fig 2A-B**). Moreover, genetic removal of *Ubx* function resulted in a loss of *GD-lacZ* repression in A1 segments, whereas *abd-A* mutant embryos maintained significant repression in all abdominal segments (**Fig 2C-E**). Thus, both Ubx and Abd-A can repress *GD-lacZ* activity in Slp2+ cells, and analysis of *GD-lacZ* reporter activity in the A1 abdominal segment specifically measures Ubx-dependent transcriptional repression.

**Figure 2.**
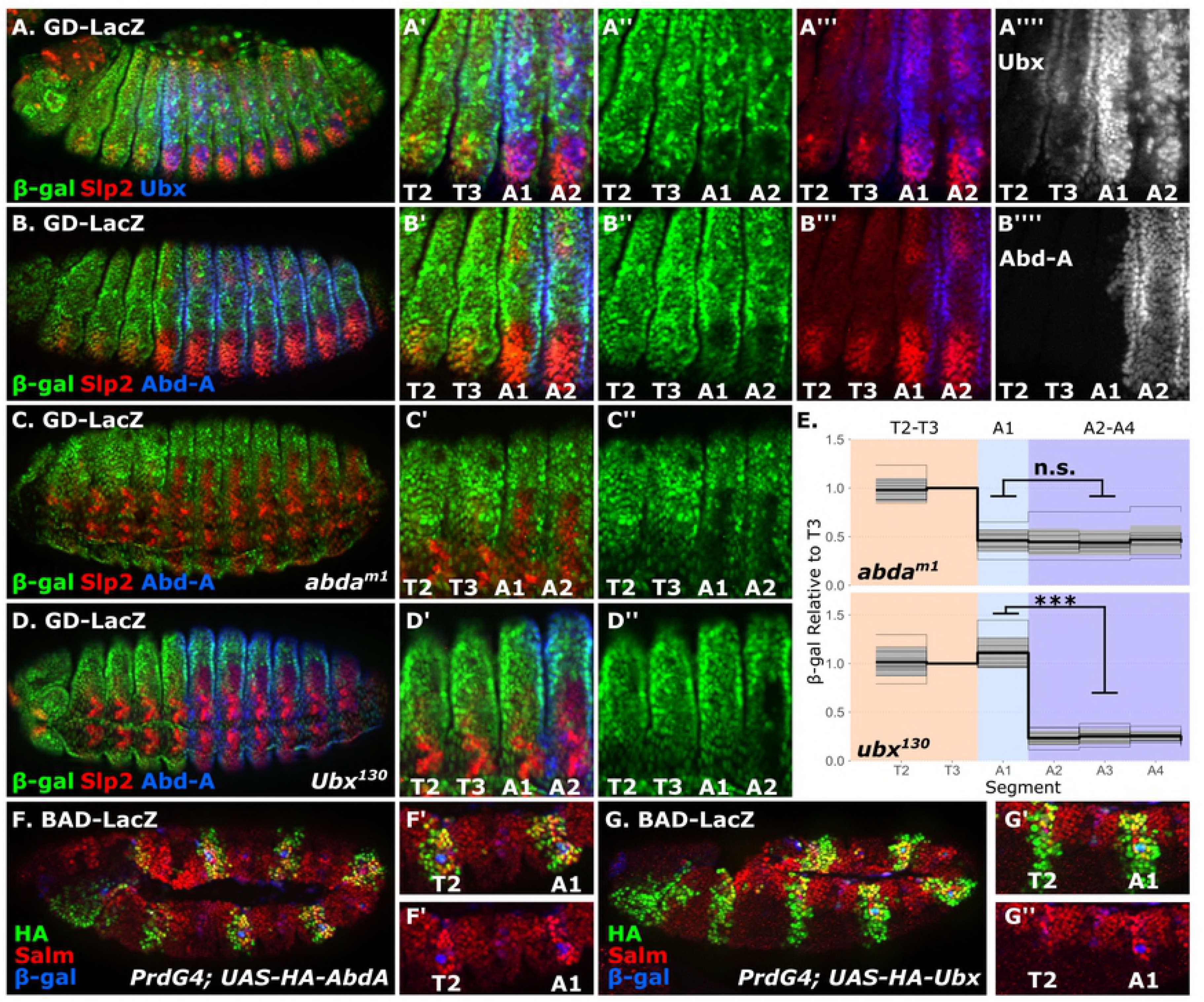
Abd-A and Ubx both repress the *DCRE* but only Abd-A activates *RhoA*. **(A)** Lateral view of stage 15 embryo carrying the *GD-LacZ* reporter immunostained for β-gal (green), Slp2 (red), Ubx (blue). **(B-D)** Lateral view of stage 15 wildtype (B), *abdA^m1^* (C), and *Ubx^130^* mutant (D) embryos carrying the *GD-LacZ* reporter immunostained for β-gal (green), Slp2 (red), Abd-A (blue). **(E)** Quantification of β-gal immunostain intensity of *GD-LacZ* in *abdA^m1^* and *Ubx^130^* mutant backgrounds. Light gray lines are intensity values from individual embryos, while the dark black line displays median value between embryos. The grey ribbon displays mean intensity value +/- the standard deviation between embryos. Statistical comparison of β-gal intensity between A1 and mean A2-A4 abdominal segments was conducted using t-test (“n.s.” = not significant, and “***" p < 0.001). **(F-G)** Lateral view of stage 11 *PrdG4:UAS-HA-AbdA* (F) and *PrdG4;UAS-HA-Ubx* (G) embryos carrying the *RhoBAD-LacZ* reporter immunostained for HA (green), Spalt-major (Salm, red) and β-gal (blue).

To better understand the mechanisms underlying abdominal Hox outputs, we next compared the ability of Ubx and Abd-A to regulate the *RhoBAD cis*-regulatory element. Unlike *Dll* and the *DCRE*, which are repressed by abdominal Hox factors, *rhomboid* (*rho*) is activated by Abd-A in a subset of abdominal sensory organ precursor cells (SOPs) via a highly conserved Exd/Hth/Hox binding site within *RhoBAD* [25, 26]. *rho* encodes a serine protease that triggers the release of an EGF ligand and neighboring cells that receive the EGF signal are specified to form larval oenocytes [30, 31]. To determine if Ubx can activate *RhoBAD*, we used the *PrdG4* driver to ectopically express Ubx in the thorax and found that neither *RhoBAD-lacZ* nor oenocytes (marked by high Spalt-major (Salm) expression) were substantially induced in thoracic segments (**Fig 2G**). In contrast, *PrdG4;UAS-Abd-A* embryos induced both *RhoBAD-lacZ* activity and oenocytes in the thorax (**Fig 2F**). Thus, while both Abd-A and Ubx can repress the *DCRE* to inhibit leg development, only Abd-A activates *RhoBAD* to induce abdominal oenocyte cells.

A notable difference between the *DCRE* and *RhoA* sequences is the organization of the Hox, Exd, and Hth sites. *RhoA* contains a single set of contiguous Exd/Hth/Hox sites, whereas the *DCRE* has multiple Hox sites that are each coupled to either an adjacent Exd or Hth binding site (**Fig 3A**). To determine if the organization of Hox, Exd, and Hth sites contributes to Hox specificity (Abd-A and not Ubx regulation via *RhoA* binding sites), we generated transgenic *GD-lacZ* lines in which the core Exd/Hth/Hox sites of *RhoA* replaced the Hox/Exd-Hth/Hox sequences within the *DCRE* (**Fig 3B**). Intriguingly, we found that the *RhoA* Exd/Hth/Hox sites can function in the *DCRE* to repress gene expression, but only in one orientation. For example, in the arbitrarily assigned forward direction (AF) the Exd/Hth/Hox sites failed to repress whereas in the reverse orientation (AR) these same sequences mediated significant repression (**Fig 3E-F**). Importantly, electromobility shift assays (EMSAs) using purified Exd/Hth heterodimers and Abd-A revealed no significant differences in binding patterns using probes of the *DCRE-AF* versus the *DCRE-AR*, suggesting that differences in Hox DNA binding activity and complex formation with Exd/Hth cannot explain the failure of the *DCRE-AF* to mediate repression (**Fig S2**). Segment specific analysis of *GD-AR-LacZ* activity in Slp2+ cells revealed that A1 segment cells expressing only Ubx and abdominal segments expressing both Ubx and Abd-A exhibit a similar degree of repression (**Fig 3E-F**). Hence, these data show that the *RhoA* Exd/Hth/Hox sites are not strictly Abd-A specific and Ubx can utilize this configuration of sites to mediate repression when placed into the *DCRE*.

**Figure 3.**
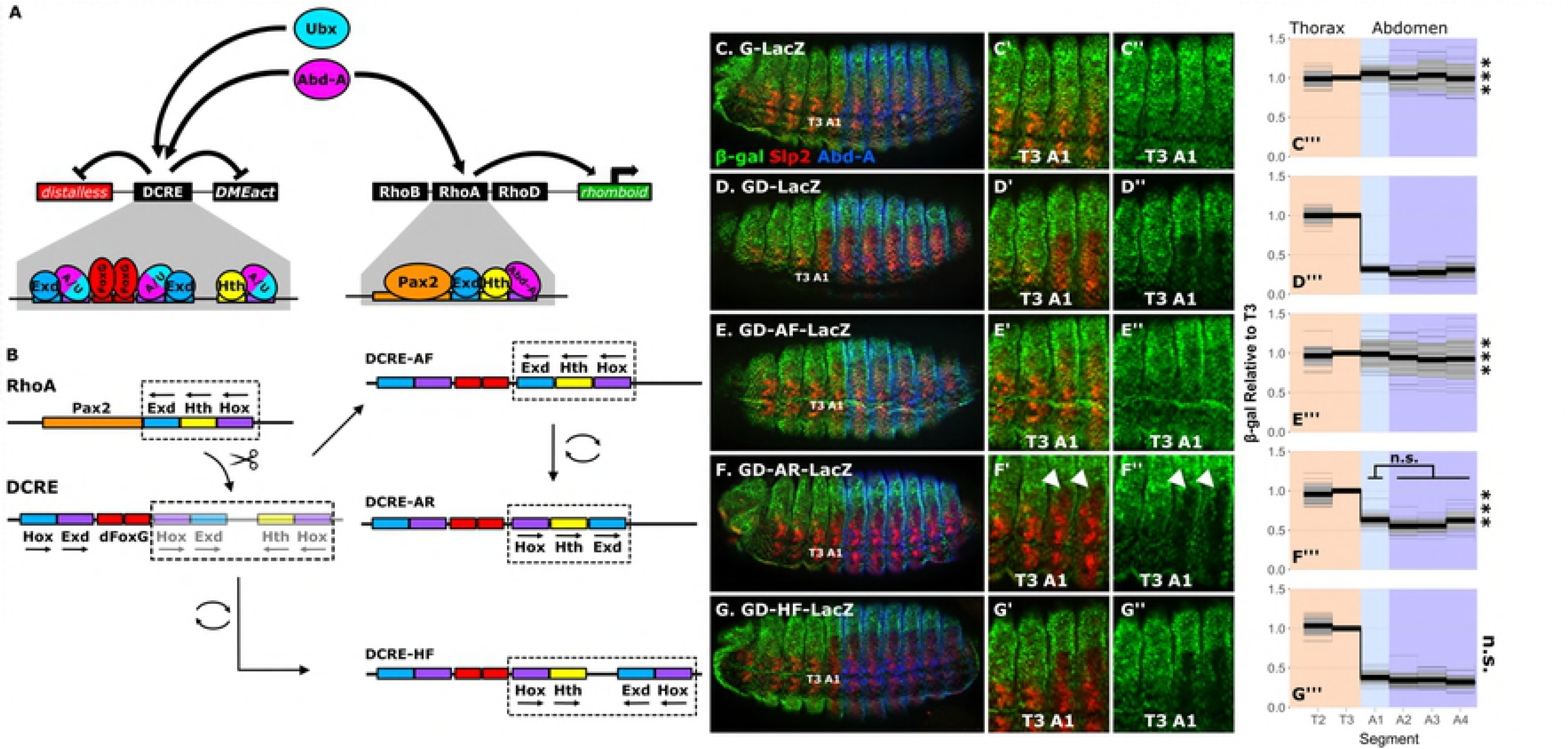
Hox, Exd, and Hth binding site arrangement does not define Abd-A vs Ubx specificity on the *RhoA* and *DCRE* elements. **(A)** A schematic of Ubx and Abd-A action on the *DCRE* and *RhoA* elements. The *DCRE* contains binding sites that recruit both Ubx and Abd-A Hox factor (A/U), Exd, Hth, and FoxG (Slp1/2) to repress transcription. The *RhoA* contains binding sites that recruit only the Abd-A Hox factor along with Exd, Hth, and Pax2 to activate transcription in abdominal SOP cells. **(B)** Schematic showing how the Hox/Exd-Hth/Hox sites in *DCRE* were replaced with the *RhoA* Exd/Hth/Hox sites to create the *DCRE-AF* and *DCRE-AR* constructs. Additionally, the Hox/Exd-Hth/Hox sites were placed in a reverse complement order to produce the *DCRE-HF* construct. **(C-G)** Lateral view of *Drosophila* embryos (stage 15) carrying the indicated *LacZ* reporter constructs immunostained for β-gal (green), Slp2 (red), and Abd-A (blue). **(C’-G’)** High resolution view of the T2-A2 segments of panels C-G. **(C’’-G’’)** Same image as (C’-G’) but only showing β-gal stain. **(C’’’-G’’’)** Quantification of β-gal immunostain intensity among Slp2+ cells in segments T2-A4 relative to intensity in T3 segment. Light gray lines are intensity values from individual embryos, while the dark black line displays median value between embryos. The grey ribbon displays mean intensity value +/- the standard deviation between embryos. Statistical comparison of mean abdominal β-gal intensity between reporter constructs was conducted using ANOVA with post-hoc Tukey’s test. Asterisks indicate significant difference from *GD-LacZ* construct (“n.s.” = not significant, “*” p < 0.05, “**" p < 0.01, “***" p < 0.001).

### The orientation and spacing of Hox sites relative to FoxG sites is critical for Abdominal Hox-mediated repression of the *DCRE*

The findings that the *RhoA* Exd/Hth/Hox sites can mediate abdominal repression in Slp+ cells when inserted into the *DCRE* in only one direction (*DCRE-AR*) suggests that Hox binding site orientation relative to the FoxG sites may be a critical factor in conveying output. Consistent with this idea, sequence comparisons between the *DCRE*, *DCRE-AF*, and *DCRE-AR* revealed that in the *DCRE-AR* the *RhoA* Hox/Exd site is in a similar orientation and spacing relative to the FoxG sites as the endogenous Hox/Exd site (**Fig 3B**). In contrast, the *DCRE-AF* does not recapitulate this spacing/orientation and the Hox/Exd site is in the opposite orientation 10 nucleotides away (**Fig 3B**). Moreover, we found that when the entire Hox/Exd-Hth/Hox sequence within the *DCRE* was "flipped" over in the opposite orientation (*DCRE-HF*), it placed the Hox/Hth site in a similar orientation and spacing relative to the FoxG site as the original Hox/Exd site and repressed Slp2+ abdominal gene expression as well as the wild type *DCRE* (**Fig 3G**).

To further test the importance of spacing between the *DCRE* FoxG and Hox sites, we inserted short DNA sequences between these sites. Care was used to ensure the inserted sequences did not code for any additional Hox, Exd, Hth, or FoxG binding sites (**Fig 4A**). Five nucleotide intervals were used to systematically alter the DNA phasing of the binding sites along the alpha helix (10 nucleotides = ~1 turn of the DNA helix). Intriguingly, we found that inserting +5 nucleotides (a half phase of the DNA helix) resulted in a complete loss of repression even though all three Hox/Hox co-factor sites and the FoxG sites are present (compare **Fig 4B** with **4C**, quantified in **4G**). In contrast, inserting +10 nucleotides partially rescued abdominal repression in Slp2+ cells, and *GD-lacZ* reporters with +15 or +20 nucleotide insertions were also able to mediate repression as well or even better than the +10 spacer (**Fig 4D-4G**). Since the +5bp insertion resulted in a complete loss of repression, we used EMSA analysis to compare DNA binding activity to wild type *DCRE*, *DCRE*+*5* and *DCRE*+*10* probes and found no significant difference in Abd-A and Exd/Hth binding (**Fig S2**). These findings indicate that the repression activity mediated by the FoxG and Hox factors is constrained by the close proximity of their binding sites within the native *DCRE* element. In contrast, when the FoxG and Hox sites are farther apart, they have added flexibility in mediating transcriptional repression.

**Figure 4.**
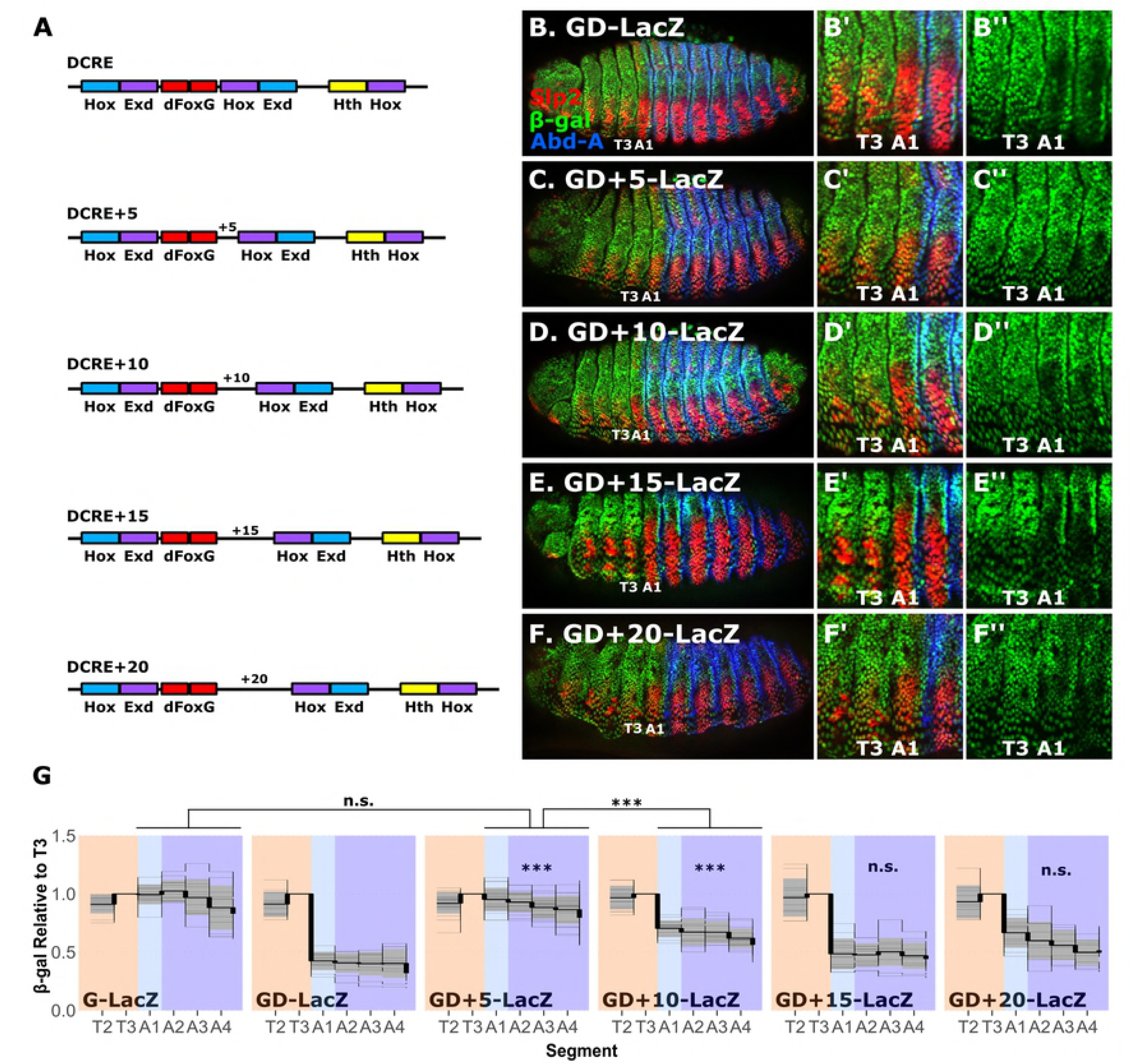
Spacing between Hox and dFoxG sites is critical for Hox-mediated repression of the *DCRE*. **(A)** Schematics of *DCRE* variants with 5, 10, 15, or 20 bp insertions between the FoxG and Hox binding sites. **(B-F)** Lateral view of *Drosophila* embryos (stage 15) carrying the indicated *LacZ* reporter constructs immunostained for β-gal (green), Slp2 (red), and Abd-A (blue). **(B’-F’)** High resolution view of T2-A2 segments of panels B-F. **(B’’-F’’)** Same as (B’-F’) but only showing β-gal stain**. (G)** Quantification of β-gal immunostain intensity of Slp2+ cells in segments T2-A4 relative to intensity in T3 segment of the indicated *LacZ* reporter embryos. Light gray lines are intensity values from individual embryos, while the dark black line displays median value between embryos. The grey ribbon displays mean intensity value +/- the standard deviation between embryos. Statistical comparison of mean abdominal β-gal intensity between reporter constructs was conducted using ANOVA with post-hoc Tukey’s test. Asterisks indicate significant difference from GD-LacZ construct (“n.s.” = not significant, “*” p < 0.05, “**" p < 0.01, “***" p < 0.001).

### The orientation of the FoxG sites is critical for Abdominal Hox-mediated repression of the *DCRE*

The FoxG (Slp) factors have been shown to directly repress gene expression via two other *cis*-regulatory elements: an *even-skipped* (*eve*) enhancer in the early *Drosophila* ectoderm and a *bagpipe* (*bap*) enhancer in the embryonic visceral mesoderm [41, 42]. Sequence comparisons between the *DCRE*, *eve*, and *bap* elements reveals each contains at least two FoxG binding sites, but in distinct orientations. The *DCRE* FoxG sites are in a head-to-head (HH) orientation, the *eve* FoxG sites are in a head-to-tail (HT) orientation, and the *bap* FoxG sites are in a tail-to-tail (TT) orientation (**Fig 5A**). To determine if the orientation of the FoxG sites is critical for *DCRE* mediated repression, we replaced the native FoxG sites with those from *bap* and *eve* (**Fig 5A**). Comparative EMSAs using *DCRE* probes with either the *bap* or *eve* FoxG sites revealed purified Slp1 protein binds both sequences as well or better than the wild type FoxG sites (**Fig 5B**). In contrast, point mutations within the wild type FoxG sites (SlpM) weakens Slp1 binding to the *DCRE* (**Fig 5B**). Intriguingly, expression analysis of *GD-lacZ* embryos containing either the *eve* or *bap* FoxG sites revealed a significant loss of repression that was comparable to the *GD-SlpM-lacZ* embryos (**Fig 5C-E** and **5H**). Similar results were also seen when the FoxG sites from the *eve* enhancer were inserted in the reverse complement orientation of the *DCRE* relative to the Hox sites (EveRC, **Fig 5F** and **5H**). To further ascertain if the orientation of FoxG sites is critical for *DCRE*-mediated repression, we re-engineered the *bap* FoxG sites into a Head-to-Head orientation within the *DCRE* (BapC) and found that it repressed as well as the wild type *DCRE* (**Fig 5G-H**). In all cases, similar behaviors were observed in the A1 segment that only expresses Ubx as compared to abdominal segments that express both Ubx and Abd-A. Taken together, these findings demonstrate that the orientation and spacing of the Hox and FoxG sites are critical for mediating robust abdominal transcriptional repression by Ubx and Abd-A in Slp+ cells.

**Figure 5.**
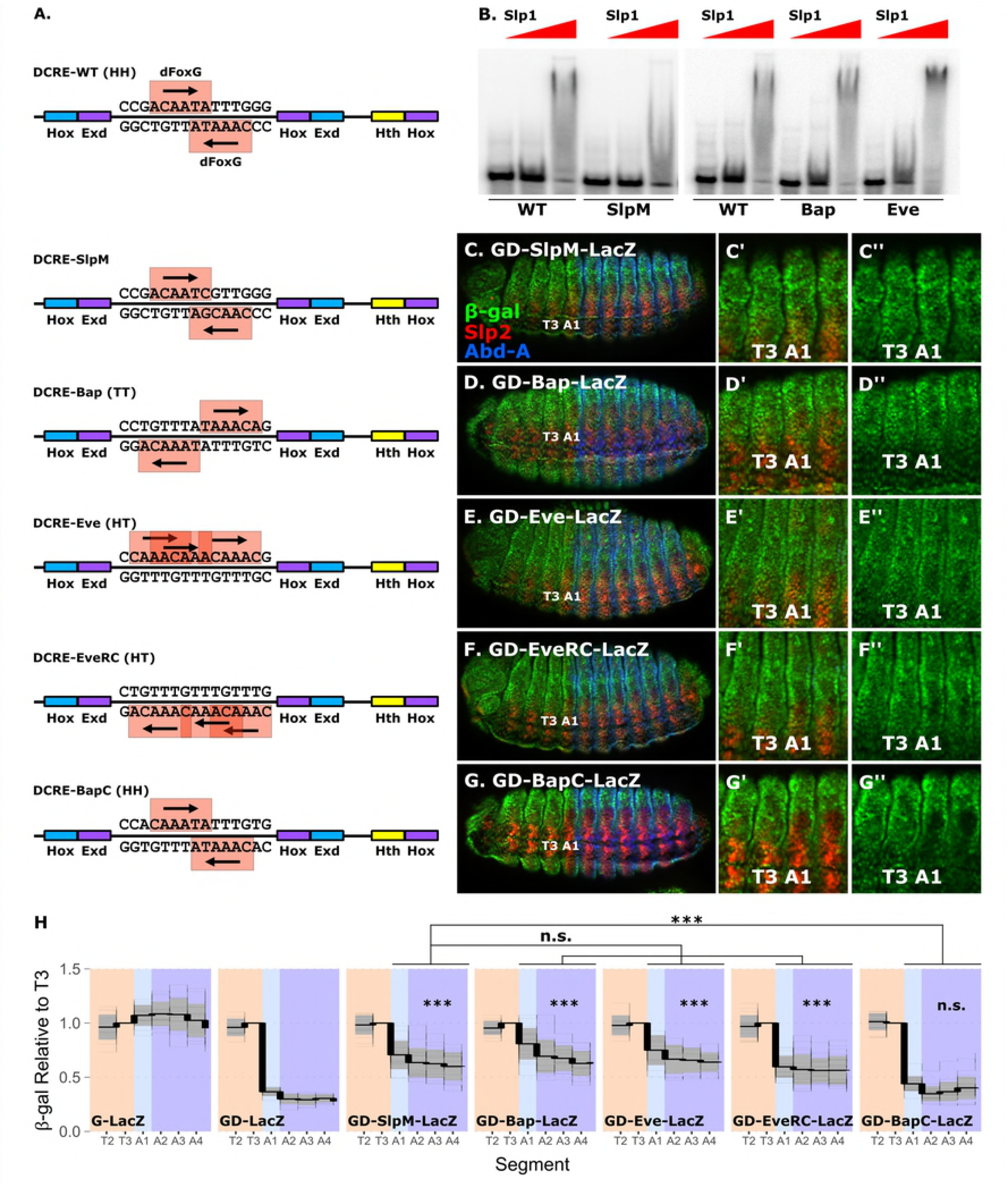
Orientation of FoxG sites is critical for Hox-mediated repression of the *DCRE*. **(A)** Schematics of *DCRE* variants: wildtype (*DCRE-WT*), weakened FoxG sites (*DCRE-SlpM*), or differing orientations of FoxG sites (*DCRE-Eve*, *DCRE-Bap*, *DCRE-Eve*). **(B)** EMSAs using two concentrations of Slp1 (100 and 500ngs) with *DCRE-WT*, *DCRE-SlpM, DCRE-Eve,* and *DCRE-Bap* probes. Note, that the SlpM sequence change disrupts most but not all Slp1 binding whereas the Bap and Eve probes bind as well or better than the wild type DCRE sequence. **(C-G)** Lateral view of Drosophila embryos (Stage 15) carrying the indicated *LacZ* reporter constructs immunostained for β-gal (green), Slp2 (red), and Abd-A (blue). **(C’-G’)** High resolution view of T2-A2 segments from panels C-G. **(C’’-G’’)** Same as (C’-G’) but only showing β-gal stain. **(H)** Quantification of β-gal immunostain intensity among Slp2+ cells in segments T2-A4 relative to intensity in T3 segment of the indicated *LacZ* reporter embryos. Light gray lines are intensity values from individual embryos, while the dark black line displays median value between embryos. The grey ribbon displays mean intensity value +/- the standard deviation between embryos. Statistical comparison of mean abdominal β-gal intensity between reporter constructs was conducted using ANOVA with post-hoc Tukey’s test. Asterisks indicate significant difference from GD-LacZ construct (“n.s.” = not significant, “*” p < 0.05, “**" p < 0.01, “***" p < 0.001).

### The spacing of Hox sites relative to Pax2 sites is critical for Abdominal Hox-mediated activation of the *RhoA* element

Our previous studies revealed that the three Hox/Hox co-factor sites within the *DCRE* can recruit abdominal Hox complexes to mediate repression [35]. In contrast, the *RhoA* element contains a single contiguous set of Exd/Hth/Hox sites that mediate activation in conjunction with a nearby Pax2 site (**Fig 3A**). To determine if the Hox/Hox co-factor sites from the *DCRE* can similarly mediate *RhoA* activation in abdominal SOP cells, we used a previously established transgenic reporter assay based on three copies of the *RhoA* element (*RhoAAA-lacZ*) which is sufficient to mediate activation in abdominal SOP cells [26, 28] (**Fig 6B**). To do so, we made *RhoAAA-DF* (forward) and *RhoAAA-DR* (reverse) constructs in which the Pax2 site was maintained but the *RhoA* Exd/Hth/Hox sites were replaced with the *DCRE* Hox/Exd-Hth/Hox sites in the "forward" and "reverse" orientations (**Fig 6A**). Interestingly, neither *RhoAAA-DR-lacZ* nor *RhoAAA-DF-lacZ* were capable of activating transcription in abdominal SOPs (**Fig 6C-D**). Given that inserting the *DCRE* Hox/Exd-Hth/Hox sites into *RhoA* alters the spacing between Hox and Pax2 sites, we next tested how spacing between the Pax2 and *RhoA* Exd/Hth/Hox sites affects Abd-A mediated activation. For this purpose, we inserted +5 or +10 nucleotide sequences between the Pax2 and Exd/Hth/Hox sites (**Fig 6E**). Since *RhoA* also encodes an overlapping Senseless (Sens) binding site that can repress thoracic gene expression, care was taken to ensure that a low-affinity Sens site was maintained and that no new Pax2, Exd, Hth, or Hox sites were created within the *RhoA* element [43]. Like the *DCRE*, we found that insertion of 5 bp sequences between the Pax2 and Exd/Hth/Hox sites resulted in a loss of Hox mediated activation. However, unlike the *DCRE*, the *RhoA* element did not regain any activity when a full helical phase of DNA sequence (+10 bp) was inserted between these sites. Thus, these findings are consistent with the Pax2 and Exd/Hth/Hox sites being highly constrained in order to mediate transcriptional activation in abdominal SOPs.

**Figure 6.**
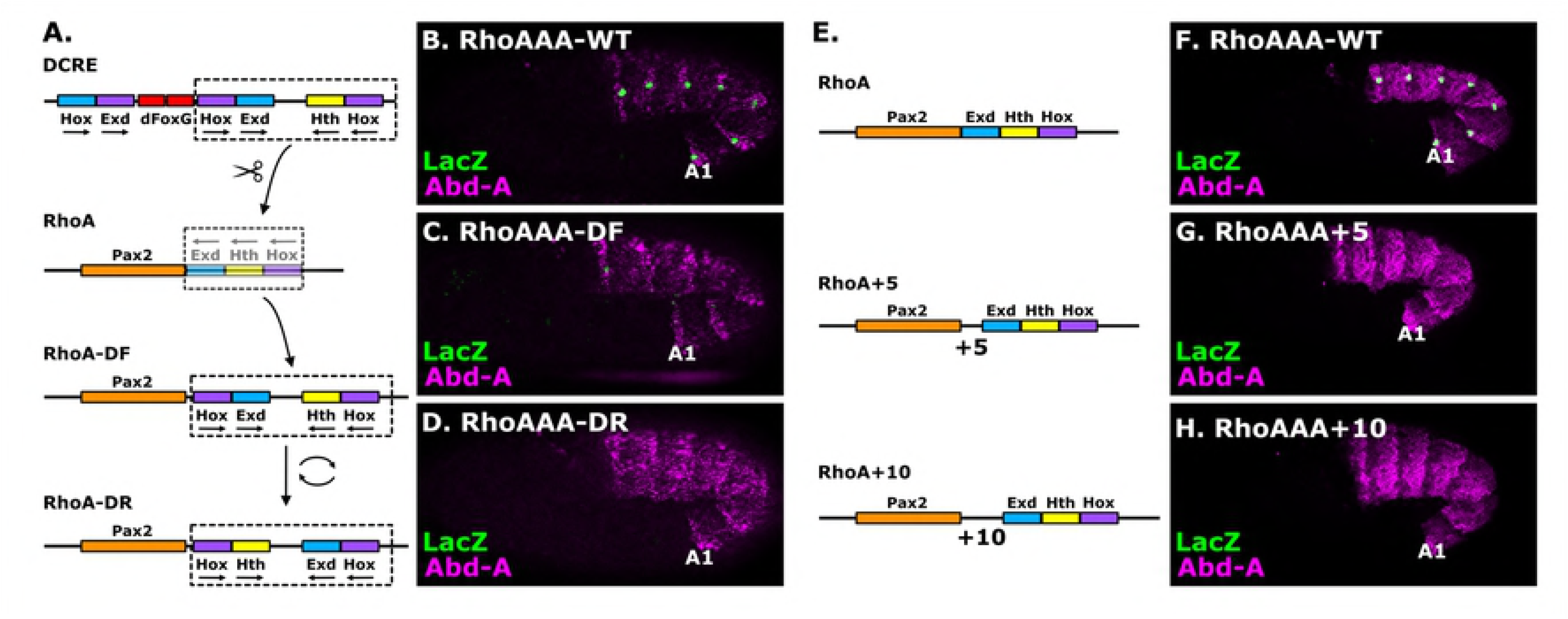
The arrangement and spacing of the Hox, Exd, and Hth binding sites are critical for *RhoA* activity. **(A)** Schematic demonstrating how the Exd/Hth/Hox binding sites in *RhoA* were replaced with the Hox/Exd-Hth/Hox binding sites from the *DCRE* to create the *RhoA-DF* and *RhoA-DR* constructs. **(B-D)** Lateral view of *Drosophila* embryos (stage 10) carrying the indicated *LacZ* reporter construct immunostained for β-gal (green), and Abd-A (magenta). **(E)** Schematic of *RhoA* variants with an additional 5 or 10 bp between the Pax2 and Exd-Hth-Hox binding sites. **(F-H)** Lateral view of *Drosophila* embryos (stage 10) carrying the indicated *LacZ* reporter construct immunostained for β-gal (green), and Abd-A (magenta).

## Discussion

A long-standing question in developmental biology has been how homologous transcription factors with similar DNA binding domains produce different activities. The Hox homeodomain factors are exemplars of such protein families, as each Hox factor binds similar AT-rich sequences *in vitro*, yet drives distinct developmental programs *in vivo* [7]. One advance in answering this paradox was to show that Hox proteins have different binding preferences in the presence of Exd and Hth [11-13]. Examples include: posterior Hox factors have greater affinity for adjacent Hth/Meis sites than anterior Hox factors [44, 45], and Hox interactions with Exd/Pbx proteins via the Hox hexapeptide motif (YPWM) can uncover latent DNA binding specificity that better discriminates between Hox factors [19]. More recently, subsets of Hox factors have been shown to mediate additional interactions with Pbx/Exd proteins via specific PBC-interaction motifs (SPIMs) [14, 15]. While SPIM interactions between Hox and PBC proteins are thought to be weaker and more dynamic than those mediated by the classic YPWM motif, it is possible they further aid in the ability of Hox factors to bind distinct DNA sequences. Thus, these findings suggest that distinct modes of interactions between Hox and Exd/Pbx and Hth/Meis proteins can alter complex formation on DNA and thereby affect target specificity.

In this study, we investigated the mechanisms underlying how the Hox, Exd, and Hth binding sites of the *RhoA* and *DCRE* regulatory elements mediate specific transcriptional outcomes. First, we found that the *DCRE* mediates transcriptional repression over a relatively short range as moving it ~700bps from activator sequences abolishes its ability to repress transcription. This finding is consistent with published ChIP-PCR data showing that Ubx and Abd-A repress *dll* expression by selectively binding to the distal DCRE and not the *dll* promoter *in vivo* [37]. Second, we found that the *RhoA* Exd/Hth/Hox sites, which mediate Abd-A specific activation in abdominal SOP cells, can also be used by the Ubx Hox factor to mediate repression when inserted into the *DCRE*. Hence, the *RhoA* configuration of Exd/Hth/Hox sites are not strictly Abd-A specific binding sites. Third, since Abd-A can use the same set of Exd/Hth/Hox sites to mediate activation in the *RhoA* element and repression in the *DCRE*, these data show that this configuration of binding sites does not confer activation vs repression. Instead, the integration of Exd/Hth/Hox complexes with nearby transcription inputs are required to discriminate between activation vs repression. Furthermore, only Abd-A, and not Ubx, is capable of mediating activation when bound to the *RhoA* element. Hence, the ability of Ubx and Abd-A to accurately regulate target genes and ultimately distinct cell fates is unlikely to solely rely upon differences in DNA binding affinity mediated by the Exd and Hth Hox co-factor proteins. Altogether, these findings reveal insights into the mechanisms used to ensure accurate Hox-specific target regulation and into the grammar underlying how *cis*-regulatory modules yield cell- and segment-specific outputs.

### Hox specificity: Target activation vs target repression

What defines whether a Hox/Exd/Hth complex activates or represses transcription once bound to DNA? Our comparative studies of the *DCRE* and *RhoA* elements show that the presence of additional binding sites for other transcription factors is critical to mediate appropriate output. For example, in addition to an Abd-A/Hth/Exd Hox complex, *RhoA* requires an appropriately positioned Pax2 binding site to mediate abdominal SOP gene activation. Since Pax2 is not expressed exclusively in the abdomen but is expressed in all *Drosophila* segments, these findings suggest that Pax2 selectively works with abdominal Hox factors via a nearby DNA binding site. Consistent with this idea, previous studies demonstrated that Abd-A and Pax2 could be co-immunoprecipitated in cell culture, whereas a thoracic Hox factor (Antennapedia, Antp) that fails to activate *RhoA* also failed to form such complexes with Pax2 [26]. Moreover, the vertebrate Hox11 proteins were also found to form protein complexes with Pax2 to regulate target gene expression in the mammalian kidney [46]. These studies suggest that abdominal Hox/Pax2 interactions are a conserved mechanism to regulate target gene expression in a tissue- and segment-specific manner.

In contrast to the *RhoA* element that mediates Abd-A specific transcriptional activation with Pax2, the *DCRE* contains two binding sites for the FoxG (Slp) transcription factors and the spacing and orientation of the FoxG and Hox binding sites is critical to mediate abdominal transcriptional repression by the Ubx and Abd-A Hox factors. Like Pax2, the FoxG factors are expressed in all *Drosophila* segments and yet regulate the *DCRE* in only abdominal segments that express Ubx and/or Abd-A [40]. While less is known about how Hox factors interact with Slp/FoxG, recent bimolecular-fluorescence (BiFC) assays in *Drosophila* found that Slp2 interacts with both Abd-A and Ubx in embryos, and at least Abd-A does so in a manner dependent on its ability to bind DNA [47]. Moreover, the thoracic Hox factor, Antp, failed to interact with Slp2 in BiFC assays, and we previously found that instead of mediating repression Antp stimulates the *DMX* leg enhancer in a *DCRE*-dependent manner via unknown mechanisms [35]. Altogether, these findings are consistent with Slp2 selectively working with Ubx and Abd-A on the *DCRE* to mediate abdominal repression. However, it should also be noted that of the five different Hox factors tested in BiFC assays, only Antp failed to interact significantly with Slp2 [47]. Thus, it is possible that the Slp/FoxG factors are directly integrated with several Hox factors and that the specificity of output will depend upon the presence of appropriately spaced/oriented DNA binding sites within the *cis*-regulatory modules.

### *cis*-regulatory grammar: The role of orientation and spacing between Hox and Pax2/FoxG binding sites in mediating proper transcriptional output

*cis*-regulatory modules (CRMs) integrate diverse transcriptional inputs to mediate cell-specific output. Studies over the past 20 years have begun to focus on how transcription factor binding sites are organized to yield appropriate output. Current models of CRM function include the flexible billboard, the enhanceosome, and the transcription factor (TF) collective [48]. As its name implies, the flexible billboard simply requires binding sites to be present within the CRM and their spacing/orientation has little effect on transcriptional output [49]. Hence, transcription output is largely additive and the loss of any one binding site often has only a modest impact on overall transcription levels. In contrast, the enhanceosome requires precisely spaced and oriented sites to mediate cooperative complex formation, and the loss of any one site can disrupt both complex formation and synergistic output [50]. Lastly, the TF collective model also stipulates cooperative complex formation on DNA, but the arrangement of transcription factor binding sites needed to mediate cooperativity isn’t highly constrained because protein-protein interactions between transcription factors can compensate for changes in DNA sequence [51]. Below, we highlight how our current understanding of the integration of transcriptional inputs by the *DCRE* and *RhoA* regulatory elements reveals aspects consistent with each of these CRM models.

Our previous studies on the *RhoA* element revealed that five transcription factor inputs impact output: the Pax2 and Exd/Hth/Hox sites promote activation whereas an overlapping binding site for the Sens transcription factor represses gene expression in thoracic segments [25, 26]. More recently, we defined two key transcription factor binding site properties required for proper abdominal SOP output: First, both the Pax2 and Sens binding sites are required to be low affinity to yield cell- and segment-specific output. For example, creating a high affinity Pax2 site resulted in ectopic *RhoA* activity in additional abdominal SOP cells, whereas a high affinity Sens site resulted in *RhoA* repression in all SOP cells [43]. Second, uncoupling the Sens and Pax2 sites so that they no longer overlap and compete for binding also disrupted appropriate segment-specific *RhoA* output [43]. Here, we further show that changing the spacing and/or orientation of the Pax2 site relative to the Exd/Hth/Hox sites disrupts activity in abdominal SOP cells. Taken together, these findings suggest that the composition of the *RhoA* binding sites is highly constrained and is largely consistent with an enhanceosome-like activity. This idea is further supported by the high degree of *RhoA* sequence conservation observed across numerous *Drosophilid* species in terms of both actual binding site sequence and organization [43]. However, it should be pointed out that while Pax2 and Abd-A can interact in cells, no cooperativity has been detected between these factors on the *RhoA* DNA element [26]. In addition, since all of the tested *RhoA* manipulations failed to reconstitute transcriptional activation in abdominal SOP cells, it is possible that the introduced sequence changes disrupt additional, unknown binding sites within the *RhoA* element required to mediate transcriptional activation.

Comparable studies on the *DCRE* revealed a mixture of flexible and constrained DNA binding site features between the Hox and FoxG binding sites. First, we analyzed the ability of different Hox/Hox co-factor binding sites to mediate transcriptional repression within the *DCRE*. Prior studies revealed that the *DCRE* encodes two Hox/Exd sites and one Hox/Hth site that contribute to abdominal Hox mediated repression via a mechanism consistent with the TF collective model of CRM function [35]. For example, point mutations in any one Hox or Exd/Hth site had only a modest effect on both DNA binding and transcriptional repression. Moreover, a Hth isoform that completely lacks a DNA binding domain can still mediate cooperative abdominal Hox complex formation and repression of the *DCRE*, indicating that multiple Hox TF complexes can yield functional repression activity [52]. Here, we show that the Exd/Hth/Hox sites from the *RhoA* element can mediate cooperative complex formation and robust transcriptional repression within the context of the *DCRE*, but only in one orientation, even though both orientations of binding sites recruit similar abdominal Hox/Exd/Hth complexes *in vitro*. Hence, while multiple Hox/Exd/Hth site configurations can yield transcriptional repression in Slp+ abdominal cells, these new findings indicate unappreciated constraints exist in order to form functional TF complexes on the *DCRE*.

Second, we analyzed the role of FoxG binding site orientation in mediating transcriptional repression and found that FoxG sites within the *DCRE* work best in a head-to-head orientation over either a tail-to-tail site from a functional *bagpipe* (*bap*) enhancer or a head-to-tail orientation from a functional *even-skipped* (*eve*) enhancer [41, 42]. Moreover, adding a five base-pair spacer (+5) between the FoxG and Hox sites disrupted all repression activity, while adding longer spacer sequences (+10, +15, and +20) between these sites resulted in abdominal repression. These findings are consistent with the idea that when FoxG and Hox sites are in close proximity, such as in the wild type *DCRE* sequence, they require a precise spacing to mediate robust repression. In contrast, when the binding sites are spaced further apart they can repress transcription via a mechanism relatively insensitive to the phasing between binding sites (i.e. +15 and +20 have similar repression activity).

In sum, these findings reveal previously unknown constraints in the spacing and orientation of the *RhoA* and *DCRE* binding sites. Two possible mechanisms could explain why transcription factor binding sites are highly constrained: First, interactions between the abdominal Hox/Hth/Exd complexes and FoxG and/or Pax2 could enhance the DNA binding affinity and/or selectivity of the functional transcriptional complexes. While current studies did not show added cooperativity in DNA binding to either the *RhoA* or *DCRE* sequences *in vitro*, these studies were performed using non-full-length proteins produced in bacteria and thus lack post-translational modifications [26, 35]. Hence, additional domains and/or post-translational modifications may be necessary for the formation of cooperative transcriptional complexes. Second, the binding of different transcription factors in close proximity may recruit additional co-factor proteins required for transcriptional activation or repression. For example, an abdominal Hox complex and either Pax2 or the FoxG transcription factors may form an interaction surface necessary to recruit a transcriptional co-factor protein. Future studies are needed to fully define the molecular mechanisms underlying the integration of the segment-specific abdominal Hox/Exd/Hth complexes and the tissue-specific FoxG (Slp1/2) and Pax2 transcription factors.

## Materials and Methods

### Transgenic Reporter Assays

Oligonucletoides for *DCRE* sequence variants were ordered from Integrated DNA Technologies and cloned into the *pAttB-LacZ* plasmid containing *3xGBE*. RhoAAA sequences were similarly ordered and cloned into the *pAttB-LacZ* plasmid. DNA sequences for each site are found in Supplemental Data. The DNA spacer sequence was generated by PCR amplification of a portion of the *kanamycin* gene as previously described [39]. All plasmids were sequence confirmed prior to injection. Transgenic flies were created using the ϕ-C31 system with each construct inserted into the same locus (51C) [53]. Injections were conducted by Rainbow Transgenics Inc.

*Drosophila* stocks carrying *lacZ* transgenes were made homozygous for reporter constructs, and embryos were collected and stained using standard procedures at 25°C. The *UAS-HA-Ubx* and *UAS-HA-AbdA* lines were a kind gift from Richard Mann, and the *PrdG4;UAS-HA-Ubx* and *PrdG4;UAS-HA-AbdA* experiments were performed at 25°C. Embryos were immunostained using the following primary antibodies: chicken anti-β-gal (1:1000) (Abcam), guinea-pig anti-Abd-A (1:500) [26], rat anti-Slp2 (1:500) [35], mouse anti-Ubx (DSHB, 1:50), rabbit anti-Salm (1:2000) [54], and rat anti-HA (1:1000). Immunostains were detected using fluorescent secondary antibodies (Jackson Immunoresearch Inc and AlexaFluor). All images were taken using the Zeiss Axiocam with optical sectioning. Reporter β-gal pixel intensity was measured manually using NIH ImageJ and normalized to β-gal intensity in the T3 segment. Graphs and statistical analysis of reporter quantifications were conducted in R as described in figure legends.

### EMSAs

A His-tagged Slp1 construct was made by cloning the N-terminus through the DNA binding domain of Slp1 (amino acids #1 through 216) into a modified pET14b vector. His-tagged Exd-Hth heterodimers, Abd-A, and Slp1 protein were purified from BL21 using Ni+ beads, as described previously [34, 45]. SDS-PAGE and Coomassie blue staining was used to confirm the purification of the protein of interest. For EMSA, fluorescent DNA probes (Integrated DNA Technologies) were mixed with combinations of purified protein as indicated in Figures, incubated for 10 min prior to running on polyacrylamide gels, and imaged using an Odyssey LiCOR cLX scanner as previously described [35, 55].

## Supplemental Data and Figure Captions

**Supplemental Data: Sequences used in the GD-lacZ and *RhoAAA-lacZ* reporter assays.** Sequences in FASTA format for all the constructs generated and tested in the transgenic reporter assays. Note, the *RhoAAA-lacZ* transgenic reporters contain three concatemers of each listed *RhoA* sequence

**Figure S1. The *spacer (sp)* sequence is transcriptionally inert**. Quantification of β-gal immunostain intensity among abdominal Slp2+ cells in *G-LacZ* vs *spG-LacZ* reporter embryos.

**Figure S2. Comparative DNA binding analysis of Exd/Hth/Abd-A to *DCRE* variant sequences.** EMSA analysis of Exd, Hth, and Abd-A binding to the indicated DCRE variants. The amounts of each protein used were 59.2ngs of the Exd/Hth heterodimer and 94.5 and 189ngs of Abd-A.

## Bibliography

1. McGinnis, W. and R. Krumlauf, Homeobox genes and axial patterning. Cell, 1992. 68(2): p. 283-302.

2. Pearson, J.C., D. Lemons, and W. McGinnis, Modulating Hox gene functions during animal body patterning. Nat Rev Genet, 2005. 6(12): p. 893-904.

3. Mann, R.S., K.M. Lelli, and R. Joshi, Hox specificity unique roles for cofactors and collaborators. Curr Top Dev Biol, 2009. 88: p. 63-101.

4. Hueber, S.D., et al., Comparative analysis of Hox downstream genes in Drosophila. Development, 2007. 134(2): p. 381-92.

5. Slattery, M., et al., Genome-wide tissue-specific occupancy of the Hox protein Ultrabithorax and Hox cofactor Homothorax in Drosophila. PLoS One, 2011. 6(4): p. e14686.

6. Prasad, N., et al., A comparative genomic analysis of targets of Hox protein Ultrabithorax amongst distant insect species. Sci Rep, 2016. 6: p. 27885.

7. Zandvakili, A. and B. Gebelein, Mechanisms of Specificity for Hox Factor Activity. J Dev Biol, 2016. 4(2).

8. Noyes, M.B., et al., Analysis of homeodomain specificities allows the family-wide prediction of preferred recognition sites. Cell, 2008. 133(7): p. 1277-89.

9. Berger, M.F., et al., Variation in homeodomain DNA binding revealed by high-resolution analysis of sequence preferences. Cell, 2008. 133(7): p. 1266-76.

10. Affolter, M., M. Slattery, and R.S. Mann, A lexicon for homeodomain-DNA recognition. Cell, 2008. 133(7): p. 1133-5.

11. Moens, C.B. and L. Selleri, Hox cofactors in vertebrate development. Dev Biol, 2006. 291(2): p. 193-206.

12. Mann, R.S. and S.K. Chan, Extra specificity from extradenticle: the partnership between HOX and PBX/EXD homeodomain proteins. Trends Genet, 1996. 12(7): p. 258-62.

13. Mann, R.S. and M. Affolter, Hox proteins meet more partners. Curr Opin Genet Dev, 1998. 8(4): p. 423-9.

14. Merabet, S. and R.S. Mann, To Be Specific or Not: The Critical Relationship Between Hox And TALE Proteins. Trends Genet, 2016. 32(6): p. 334-347.

15. Ortiz-Lombardia, M., et al., Hox functional diversity: Novel insights from flexible motif folding and plastic protein interaction. Bioessays, 2017. 39(4).

16. Chan, S.K., et al., The DNA binding specificity of Ultrabithorax is modulated by cooperative interactions with extradenticle, another homeoprotein. Cell, 1994. 78(4): p. 603-15.

17. Chang, C.P., et al., Pbx modulation of Hox homeodomain amino-terminal arms establishes different DNA-binding specificities across the Hox locus. Mol Cell Biol, 1996. 16(4): p. 1734-45.

18. Chang, C.P., et al., Meis proteins are major in vivo DNA binding partners for wild-type but not chimeric Pbx proteins. Mol Cell Biol, 1997. 17(10): p. 5679-87.

19. Slattery, M., et al., Cofactor binding evokes latent differences in DNA binding specificity between Hox proteins. Cell, 2011. 147(6): p. 1270-82.

20. Joshi, R., et al., Functional specificity of a Hox protein mediated by the recognition of minor groove structure. Cell, 2007. 131(3): p. 530-43.

21. Abe, N., et al., Deconvolving the recognition of DNA shape from sequence. Cell, 2015. 161(2): p. 307-18.

22. Ryoo, H.D. and R.S. Mann, The control of trunk Hox specificity and activity by Extradenticle. Genes Dev, 1999. 13(13): p. 1704-16.

23. Joshi, R., L. Sun, and R. Mann, Dissecting the functional specificities of two Hox proteins. Genes Dev, 2010. 24(14): p. 1533-45.

24. Crocker, J., et al., Low affinity binding site clusters confer hox specificity and regulatory robustness. Cell, 2015. 160(1-2): p. 191-203.

25. Li-Kroeger, D., et al., Hox and senseless antagonism functions as a molecular switch to regulate EGF secretion in the Drosophila PNS. Dev Cell, 2008. 15(2): p. 298-308.

26. Li-Kroeger, D., T.A. Cook, and B. Gebelein, Integration of an abdominal Hox complex with Pax2 yields cell-specific EGF secretion from Drosophila sensory precursor cells. Development, 2012. 139(9): p. 1611-9.

27. Gebelein, B., The control of EGF signaling and cell fate in the Drosophila abdomen. Fly (Austin), 2008. 2(5): p. 257-258.

28. Witt, L.M., et al., Atonal, senseless, and abdominal-A regulate rhomboid enhancer activity in abdominal sensory organ precursors. Dev Biol, 2010. 344: p. 1060-1070.

29. Brodu, V., P.R. Elstob, and A.P. Gould, abdominal A specifies one cell type in Drosophila by regulating one principal target gene. Development, 2002. 129(12): p. 2957-63.

30. Elstob, P.R., V. Brodu, and A.P. Gould, spalt-dependent switching between two cell fates that are induced by the Drosophila EGF receptor. Development, 2001. 128(5): p. 723-32.

31. Rusten, T.E., et al., Spalt modifies EGFR-mediated induction of chordotonal precursors in the embryonic PNS of Drosophila promoting the development of oenocytes. Development, 2001. 128(5): p. 711-22.

32. Gutzwiller, L.M., et al., Proneural and abdominal Hox inputs synergize to promote sensory organ formation in the Drosophila abdomen. Dev Biol, 2010. 348(2): p. 231-43.

33. Gebelein, B., et al., Specificity of Distalless repression and limb primordia development by abdominal Hox proteins. Dev Cell, 2002. 3(4): p. 487-98.

34. Gebelein, B., D.J. McKay, and R.S. Mann, Direct integration of Hox and segmentation gene inputs during Drosophila development. Nature, 2004. 431(7009): p. 653-9.

35. Uhl, J.D., A. Zandvakili, and B. Gebelein, A Hox Transcription Factor Collective Binds a Highly Conserved Distal-less cis-Regulatory Module to Generate Robust Transcriptional Outcomes. PLoS Genet, 2016. 12(4): p. e1005981.

36. Vachon, G., et al., Homeotic genes of the Bithorax complex repress limb development in the abdomen of the Drosophila embryo through the target gene Distal-less. Cell, 1992. 71(3): p. 437-50.

37. Agelopoulos, M., D.J. McKay, and R.S. Mann, Developmental regulation of chromatin conformation by Hox proteins in Drosophila. Cell Rep, 2012. 1(4): p. 350-9.

38. Uv, A.E., E.J. Harrison, and S.J. Bray, Tissue-specific splicing and functions of the Drosophila transcription factor Grainyhead. Mol Cell Biol, 1997. 17(11): p. 6727-35.

39. Swanson, C.I., N.C. Evans, and S. Barolo, Structural rules and complex regulatory circuitry constrain expression of a Notch-and EGFR-regulated eye enhancer. Dev Cell, 2010. 18(3): p. 359-70.

40. Gebelein, B. and R.S. Mann, Compartmental modulation of abdominal Hox expression by engrailed and sloppy-paired patterns the fly ectoderm. Dev Biol, 2007. 308: p. 593-605.

41. Andrioli, L.P., et al., Groucho-dependent repression by sloppy-paired 1 differentially positions anterior pair-rule stripes in the Drosophila embryo. Dev Biol, 2004. 276(2): p. 541-51.

42. Lee, H.H. and M. Frasch, Nuclear integration of positive Dpp signals, antagonistic Wg inputs and mesodermal competence factors during Drosophila visceral mesoderm induction. Development, 2005. 132(6): p. 1429-42.

43. Zandvakili, A., et al., Degenerate Pax2 and Senseless binding motifs improve detection of low-affinity sites required for enhancer specificity. PLoS Genet, 2018. 14(4): p. e1007289.

44. Shen, W.F., et al., AbdB-like Hox proteins stabilize DNA binding by the Meis1 homeodomain proteins. Mol Cell Biol, 1997. 17(11): p. 6448-58.

45. Uhl, J.D., T.A. Cook, and B. Gebelein, Comparing anterior and posterior Hox complex formation reveals guidelines for predicting cis-regulatory elements. Dev Biol, 2010. 343(1-2): p. 154-66.

46. Gong, K.Q., et al., A Hox-Eya-Pax complex regulates early kidney developmental gene expression. Mol Cell Biol, 2007. 27(21): p. 7661-8.

47. Baeza, M., et al., Inhibitory activities of short linear motifs underlie Hox interactome specificity in vivo. Elife, 2015. 4.

48. Spitz, F. and E.E. Furlong, Transcription factors: from enhancer binding to developmental control. Nat Rev Genet, 2012. 13(9): p. 613-26.

49. Arnosti, D.N. and M.M. Kulkarni, Transcriptional enhancers: Intelligent enhanceosomes or flexible billboards? J Cell Biochem, 2005. 94(5): p. 890-8.

50. Merika, M. and D. Thanos, Enhanceosomes. Curr Opin Genet Dev, 2001. 11(2): p. 205-8.

51. Junion, G., et al., A transcription factor collective defines cardiac cell fate and reflects lineage history. Cell, 2012. 148(3): p. 473-86.

52. Noro, B., et al., Distinct functions of homeodomain-containing and homeodomain-less isoforms encoded by homothorax. Genes Dev, 2006. 20(12): p. 1636-50.

53. Bischof, J., et al., An optimized transgenesis system for Drosophila using germ-line-specific phiC31 integrases. Proc Natl Acad Sci U S A, 2007. 104(9): p. 3312-7.

54. Xie, B., et al., Senseless functions as a molecular switch for color photoreceptor differentiation in Drosophila. Development, 2007. 134(23): p. 4243-53.

55. Wang, G., et al., A Hox complex activates and potentiates the Epidermal Growth Factor signaling pathway to specify Drosophila oenocytes. PLoS Genet, 2017. 13(7): p. e1006910.

